# Cytotoxic and micronuclei inducing effects of petroleum ether fraction of leaf aqueous extract of *Clerodendrum viscosum* Vent. in *Allium cepa* root tip cells

**DOI:** 10.1101/2021.06.04.447064

**Authors:** Sujit Roy, Lalit Mohan Kundu, Gobinda Chandra Roy, Manabendu Barman, Sanjib Ray

## Abstract

*Clerodendrum viscosum* is a traditionally used medicinal plant. The present study aimed to analyze a detailed cytotoxic effect of the nonpolar petroleum ether fraction (AQPEF) of leaf aqueous extract of *C. viscosum* Vent. (LAECV) in *Allium cepa* root tip cells. The LAECV was fractionated with petroleum ether and tested for *A. cepa* toxicity at early hours (2 and 4 h treatment) at concentrations 0, 50, 100, 150 and 200 µg mL^-1^. The highest aberrant cell percentage (10.45±0.46%) was scored from 150 µg mL^-1^ followed by100 µg mL^-1^ (8.75±0.26%) concentration at 4 h treated samples. The AQPEF treatment induced a significant (p<0.0001) increase in micronuclei frequency; 4.31±0.33, 5.08±0.13, 5.05±0.22 and 3.05±0.37% respectively at concentrations 50, 100, 150, and 200 µg mL^-1^. The highest polyploid frequency (20.14±0.68 %) induced with 100 µg mL^-1^ of AQPEF at 16 h recovery. 150 µg mL^-1^ is the most effective concentration of AQPEF to decipher its activity similar to colchicine (150 µg mL^-1^). In summary, the present study indicates petroleum ether is suitable for extraction of the active phytochemicals of LAECV having cytotoxic effects on *A. cepa* root tip cells. The AQPEF has colchicine like micronuclei, polyploidy, and mitotic abnormality inducing potentials in *A. cepa* root apical meristem cells.

## Introduction

*Clerodendrum viscosum* (common name: bhant, ghentu, Family: Lamiaceae) is traditionally used medicinal plant. It has gained a reputation in traditional Ayurvedic, Unani, and Homeopathic medicine (Nandi *et al*. 2015). In the Indian Homeopathic system, it is prescribed to treat diarrhea, postnatal complications, and fresh wounds (Nadkarni and Nadkarni 2002, Hamilton 1997). In Unani medicine, this plant is well known for its effectiveness against rheumatism (Singh *et al*. 1997). To treat worm infection, cough, asthma, itching, leprosy, scorpion sting, bronchitis, fever, *etc*., the whole plant decoction is used (Kirtikar and Basu 1991, Bhattacharya *et al*. 1981). The Indian Tribals of Chotanagpur plateau use this plant parts’ extracts as a remedy for malaria, cataract, diseases of skin, blood, and lung (Kirtikar and Basu 1991). The leaf methanolic extract exerts blood glucose level restoring effects after streptozotocin treatment (Das *et al*. 2011, Arvind *et al*. 2002). The leaf extract showed peripheral analgesic activity at a dose of 200 mg/kg body and a dose 500 mg/kg body weight reduced blood glucose level from 130 to 36 mg/dL in 2 h (Hossain *et al*. 2014, Chandrashekar *et al*. 2012). Allelochemicals present in the leaf aqueous extract have harmed the growth and germination of weeds in agro-ecosystem (Devi *et al*. 2013, Qasem *et al*. 2001). Leaf and stem aqueous extracts of *C. viscosum* has resulted in significant insecticidal activity against tea pests *Helopeltis theivora* and *Oligonychus coffeae* when compared to acaricide and *Azadirachta indica* cytotoxic lethality (Azad *et al*. 2013, Islam *et al*. 2013). A promising positive correlation was established between the plant’s parts and their insect repellent and insecticidal activity (Muh *et al*. 2014).

The crude leaf extracts of *C. viscosum* contain phenolics viz. fumaric acids, acetoside, methyl esters of caffeic acids, flavonoids such as apigenin, acacetin, scutellarein, quercetin, cabrubin, hispidulin, terpenoids like clerodin, steroids such as clerodone, clerodol, clerodolone, clerosterol and some fix oils containing linolenic acid, stearic acid, oleic acids, and lignoceric acid (Saroj 2016).Acute toxicity test revealed that these plant parts are safe up to 2000 mg/kg body weight (Gupta *et al*. 2012).

In our previous study, we have reported cytotoxic effects of LAECV on root apical meristem cells of wheat and onion (Ray *et al*. 2012). The metaphase arrest and cell cycle delay inducing effects were somewhat comparable to colchicine’s actions (Ray *et al*. 2017, Kundu *et al*. 2016, Ray *et al*. 2013). In another comparative study, treatment of colchicine and LAECV on *Allium cepa* root tip cells revealed their similar cytotoxic effects including an increased frequency in mitotic abnormalities and micronucleus (Kundu *et al*. 2016). Recently, Roy and Roy (2019) assessed the cytotoxic effects of aqueous and methanolic leaf extracts of *C. viscosum* treatment on *A. cepa* roots after 5 days, however, cytotoxic activities were not studied at the early hours of treatment, and also the used concentrations of the extracts were not conclusive. In continuation of the previous reports, here non-polar solvent petroleum ether was used to extract the non-polar fraction of LAECV and analyzed its cytotoxic effects at early hours, 2 to 4 h, after the AQPEF treatment (50-200 µg mL^-1^) in *A. cepa* root apical meristem cells and also at 16 h recovery. Additionally, colchicine induced cytotoxic effects were compared with the AQPEF induced cytotoxic effects on *A. cepa* root apical meristem cells.

## Materials and methods

### Plant collection and AQPEF extraction

The collected plant was taxonomically identified as *Clerodendrum viscosum* Vent. by Professor A. Mukherjee, Department of Botany, The University of Burdwan. For future reference, a voucher specimen (No. BUTBSR011) is maintained in the Department of Zoology, B.U.

Fresh leaves of *C. viscosum* were collected from Burdwan University campus, washed in tap water, dried in shade, ground by Philips Mixer Grinder HL1605, and for further use, the ground leaf was stored in a sealed container. 100 g of ground leaf was extracted in 2.5 L of boiling distilled water for 2-3 h and the extract (LAECV) was filtered with filter paper. With the help of a magnetic stirrer, the LAECV was fractionated with petroleum ether and the resulting yellow colored petroleum ether (AQPEF) solution was concentrated by rotary vacuum evaporator and stored in a glass bottle.

### Cytotoxic effects *of AQPEF* in *A. cepa* root apical meristem cells

#### Root sprouting, treatment and squash preparation

Mitotic abnormalities, micronuclei, and polyploid cells frequencies were analyzed for the elucidation of AQPEF induced cytotoxic effects on *A. cepa* root-tip cells. The similar size *A. cepa* bulbs were surface sterilized by 1% sodium hypochlorite and allowed for root sprouting. The onion bulbs were placed in 6-well plates containing distilled water and kept in dark, within an environmental test chamber at 25-27 °C. The similar-sized (2-3 cm), 48 h aged, *A. cepa* roots were treated with AQPEF (0, 50, 100, 150, and 200 µg mL^-1^) and colchicine (150 µg mL^-1^) for 2 and 4 h. After the treatment hours, 8-10 roots were fixed and processed for squash preparation following the standard procedure (Chaudhury and Ray 2015). The remaining roots were allowed to grow further for another 16 h in distilled water and subsequently, root tips were fixed. The treated and untreated root tips were fixed in aceto-methanol (3 parts methanol: 1part glacial acetic acid) for 24 h and then hydrolyzed for 10 min in 1 N HCl at 60°C. The roots were stained with 2% aceto-orcein and finally squashed in 45% acetic acid (Sharma and Sharma 1999, Ray *et al*. 2013). The well-spread areas of squashed roots were focused under the bright field light microscope for observation and scoring the mitotic abnormalities.

### Scoring and statistical analysis

In the case of squash preparation of *A. cepa* root apical meristem cells, at least three randomly coded slides were observed under the light microscope. Calculation of the mitotic index was done by counting the number of dividing cells per total cells scored for each concentration. Aberrant cell frequencies were calculated by counting the number of abnormal cells scored per slide for each concentration (Bakare *et al*. 2000). The mitotic abnormalities were analyzed by 2×2 contingency χ2-test. Pearson’s correlation matrix was formed by GraphPad Prism software 8.4.3.

## Results and discussion

The different extracts of *C. viscosum* have various pharmacological activities (Kundu *et al*. 2016, Ray *et al*. 2013, Ray *et al*. 2012, Aley *et al*. 2011, Gouthamchandra *et al*. 2010, Jirovetz *et al*. 1999, Warrier *et al*.1996, Yusuf *et al*. 1994). The present investigation analyzed a detailed cytotoxic effect of the AQPEF, a non-polar fraction of LAECV, in *A. cepa* root apical meristem cells at early hours (2 and 4 h) of treatment and also at 16 h recovery.

The cytotoxic potentials of the various phytochemical substances can be deciphered by studying the mitotic abnormalities (Caritá *et al*. 2008). A concentration-dependent increased aberrant cell percentages were observed at 2 and 4 h in AQPEF (100, 150 & 200 µg mL^-1^) treated samples. The highest aberrant cell percentage (10.45±0.46%) was scored from 150 µg mL^-1^ followed by 100 µg mL^-1^ (8.75±0.26%) concentration at 4 h treated samples. However, in 16 h recovery samples, the AQPEF induced aberrant cell frequency was considerably decreased compared to 2 and 4 h treated samples (Table 1). Colchicine (150 µg mL^-1^) induced 10.55±0.83, 8.12±0.37 and 0.50±0.17% aberrant cells respectively in 2, 4, and 4 h treatment followed by 16 h recovery samples (4 h T+16 h RS). The AQPEF induced similar aberrant cells percentage at a much lower concentration than crude aqueous extract (LAECV) at 2 and 4 h treated *A. cepa* root apical meristem cells. In the case of 4 h T+16 h RS, the aberrant cell frequencies reduced in comparison to 2 and 4 h treated cells but still aberrant cells persist more than the untreated ones. Our earlier reports indicated that LAECV has colchicine-like metaphase arrest and mitotic abnormalities inducing potentials (Kundu *et al*. 2016, Ray *et al*. 2012) and this study explored that the petroleum ether fraction (AQPEF) of LAECV contains the active principles having colchicine like aberrant cell inducing potentials.

**Table 1:** Petroleum ether fraction of leaf aqueous extract of *Clerodendrum viscosum*, AQPEF, induced micronuclei and mitotic abnormalities in *A. cepa* root apical meristem cells.

| H | Conc.<br>($\mu\text{g mL}^{-1}$) | TC | TDC | Ac | Sti | Bri | Pd | Vag | Lag | MN | POL |
| --- | --- | --- | --- | --- | --- | --- | --- | --- | --- | --- | --- |
| % (Mean±SEM) |  |  |  |  |  |  |  |  |  |  |  |
| 2 | 0 | 1959 | 111 | 0.30±0.09 | 1.78±0.89 | 1.83±1.83 | 0.00±0.00 | 0.00±0.00 | 0.00±0.00 | 0.00±0.00 | 0.00±0.00 |
|  | 50 | 2181 | 144 | 2.58±0.24<sup>a</sup> | 6.23±1.24 | 7.0±1.5 | 2.06±0.03 | 0.00±0.00 | 1.03±0.60 | 0.00±0.00 | 0.00±0.00 |
|  | 100 | 2445 | 183 | 4.59±0.33<sup>a</sup> | 5.49±1.36 | 5.62±1.08 | 5.32±1.08 | 2.67±0.35 | 2.29±0.80 | 0.00±0.00 | 0.00±0.00 |
|  | 150 | 2404 | 234 | 7.13±0.31<sup>a</sup> | 3.84±0.69 | 7.24±0.63 | 3.42±1.48 | 2.50±0.63 | 0.40±0.40 | 0.00±0.00 | 0.00±0.00 |
|  | 200 | 2389 | 251 | 6.65±0.10<sup>a</sup> | 6.42±0.52 | 7.23±0.51 | 3.54±1.24 | 0.84±0.42 | 0.81±0.41 | 0.00±0.00 | 0.00±0.00 |
|  | 150© | 1987 | 248 | 10.55±0.83<sup>a</sup> | 2.67±0.80 | 2.55±1.81 | 0.00±0.00 | 1.50±0.75 | 0.00±0.00 | 0.00±0.00 | 0.00±0.00 |
| 4 | 0 | 2914 | 207 | 0.20±0.05 | 0.00±0.00 | 3.09±1.09 | 0.00±0.00 | 0.00±0.00 | 0.00±0.00 | 0.00±0.00 | 0.00±0.00 |
|  | 50 | 3423 | 352 | 6.28±0.59<sup>a</sup> | 1.76±1.31 | 5.63±0.41 | 1.43±0.58 | 2.17±0.84 | 0.60±0.30 | 0.00±0.00 | 0.00±0.00 |
|  | 100 | 2162 | 222 | 8.75±0.26<sup>a</sup> | 0.00±0.00 | 3.60±0.47 | 0.00±0.00 | 1.35±0.05 | 0.00±0.00 | 0.00±0.00 | 0.00±0.00 |
|  | 150 | 3035 | 403 | 10.45±0.46<sup>a</sup> | 1.59±0.80 | 5.30±2.20 | 0.21±0.21 | 1.16±0.52 | 0.00±0.00 | 0.00±0.00 | 0.00±0.00 |
|  | 200 | 3525 | 352 | 7.61±0.76<sup>a</sup> | 1.66±0.76 | 5.50±0.86 | 0.35±0.35 | 0.00±0.00 | 0.00±0.00 | 0.00±0.00 | 0.00±0.00 |
|  | 150© | 3286 | 301 | 8.12±0.37<sup>a</sup> | 0.93±0.40 | 3.45±1.07 | 1.00±0.07 | 0.35±0.35 | 0.00±0.00 | 0.00±0.00 | 0.00±0.00 |
| 4+16 | 0 | 2449 | 241 | 0.25±0.03 | 0.00±0.00 | 2.49±0.10 | 0.00±0.00 | 0.00±0.00 | 0.00±0.00 | 0.00±0.00 | 0.00±0.00 |
|  | 50 | 3946 | 276 | 1.05±0.9 | 0.00±0.00 | 2.17±0.06 | 0.36±0.36 | 0.00±0.00 | 0.00±0.00 | 4.31±0.33<sup>a</sup> | 11.96±0.48<sup>a</sup> |
|  | 100 | 2130 | 177 | 4.37±0.86<sup>a</sup> | 1.16±0.63 | 3.23±1.64 | 0.59±0.59 | 3.69±0.91<sup>b</sup> | 0.00±0.00 | 5.08±0.13<sup>a</sup> | 20.14±0.68<sup>a</sup> |
|  | 150 | 2068 | 136 | 3.84±0.59<sup>a</sup> | 1.04±1.04 | 5.83±1.72<sup>c</sup> | 0.00±0.00 | 5.16±0.68<sup>a</sup> | 0.00±0.00 | 5.05±0.22<sup>a</sup> | 18.43±0.21<sup>a</sup> |
|  | 200 | 2162 | 161 | 3.13±0.52<sup>a</sup> | 4.33±1.51<sup>b</sup> | 7.38±0.71<sup>c</sup> | 0.00±0.00 | 5.04±0.83<sup>b</sup> | 0.00±0.00 | 3.05±0.37<sup>a</sup> | 12.42±0.81<sup>a</sup> |
|  | 150© | 1922 | 69 | 0.50±0.17 | 0.00±0.00 | 2.70±1.60 | 0.85±0.85 | 0.85±0.85 | 0.00±0.00 | 6.11±0.43<sup>a</sup> | 16.03±0.20<sup>a</sup> |

<sup>a</sup> significant at p<0.0001 and <sup>b</sup> significant at p<0.001, <sup>c</sup> significant at p<0.05 as compared with their respective control by 2x2 Contingency $\chi^2$ -test with respective df = 1. H; Hours, Comp; Compound, Conc; Concentration, Tc; Total number of cells, TDC; Total dividing cells, Ac; Aberrant cells, Sti; Stickiness, Bri; Bridge, Pd; Polar deviation, Vag; Vagrant chromosome, Lag; Laggard chromosome, MN; Micronucleus, POL; Polyploidy, ©; Colchicine.

The mitotic abnormalities like anaphase bridges, stickiness, polar deviation, vagrant chromosome, laggard chromosome, micronuclei, polyploidy, *etc*. were induced by both AQPEF and colchicine treatments in *A. cepa* root tip cells. The AQPEF treatment induced an increased percentage of anaphase bridges in *A. cepa* root apical meristem cells. The anaphase bridge percentages 7.0±1.5, 5.62±1.08, 7.24±0.63, and 7.23±0.51 were observed at 2 h respectively at concentrations of 50, 100, 150, and 200 µg mL^-1^ and correspondingly, these were decreased to 5.63±0.41, 3.60±0.47, 5.30±2.20, and 5.50±0.86% at 4 h and 2.17±0.06, 3.23±1.64, 5.83±1.72 and 7.38±0.71% in 4 h T+16 h RS, except for the concentration of 200 µg mL^-1^, which showed the highest (7.38±0.71%) anaphase bridge percentage (Table 1, Figure 1). Colchicine (150 µg mL^-1^) induced anaphase bridge 2.55±1.81, 3.45±1.07 and 2.70±1.60% respectively in 2, 4, and 4 h T+16 h RS. The LAECV has anaphase bridge inducing capabilities in 4 h and 4 h treatment followed by 16 h recovery *A. cepa* root tip cells (Kundu *et al*. 2016). The AQPEF at a lower concentration has produced the similar types of mitotic abnormalities, as induced in LAECV, in all the treated hours may be active principles were concentrated in the petroleum ether fraction.

**Figure 1:**
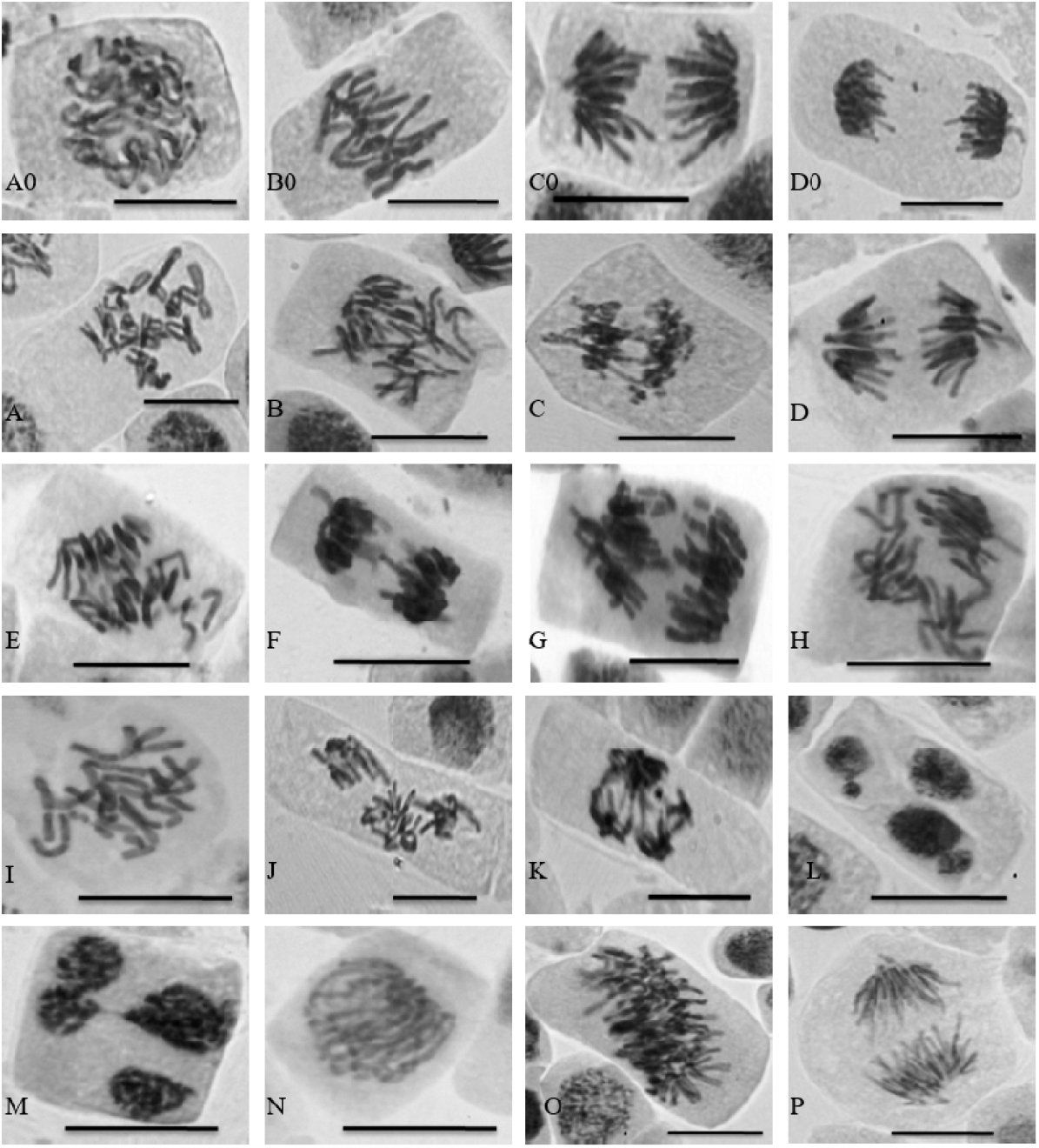
Petroleum ether fraction of leaf aqueous extract of *Clerodendrum viscosum*, AQPEF, induced polyploidy, micronuclei, and mitotic abnormalities in *A. cepa* root apical meristem cells. Untreated (A0-D0): A0; Prophase, B0; Metaphase, C0; Anaphase, and D0; Telophase of *A. cepa* root apical meristem cells. A; C-metaphase, B; Anaphase bridge, C; Decondensed anaphase bridge, D; Polar deviation, E; Vagrant chromosome, F; Sticky with anaphase bridge, G; Chromatid Break, H; Disrupted anaphase, I; Disrupted metaphase, J; Multipolar anaphase, K; Decondensed multipolar anaphase, L; Micronucleus with multiple nuclei, M; Multi nuclei with decondensed anaphase bridge, N; Polyploid prophase, O; Polyploid metaphase, P; Polyploid anaphase.

Like anaphase bridge, an increased frequency of chromosomal stickiness was also observed in both the AQPEF and colchicine treated onion root tip cells at 2 h and that was successively decreased at 4 h and 4 h T+16 h RS. The AQPEF (200 µg mL^-1^) induced the highest frequency of chromosomal stickiness (6.42±0.52%) at 2 h in onion root tip cells. In the case of 16 h recovery samples, the percentages of sticky chromosomes containing cells were reduced (Table 1, Figure 1). Similarly, colchicine (150 µg mL^-1^) induced 2.67±0.80% and 0.93±0.40% chromosomal stickiness respectively in 2 and 4 h treated samples. The basis for the onset of the sticky chromosome may be due to sub chromatid association among chromosomes or dissolution of nucleoprotein or de-condensation of DNA (Babich et al.1997, Ford and Correll 1992). The AQPEF induced anaphase bridges are may be due to fission and fusion of the chromatids and chromosomes. The AQPEF induced multipolar anaphase in condensed and decondensed form and such multipolar anaphases might be formed from the chromosomal bridge and sticky chromosome.

The percentage of polar deviation increased in onion root apical meristem cells that were treated with AQPEF, at 2 h but decreased at 4 h treatment and 4 h T+16 h RS. At 2 h, AQPEF induced polar deviation were scored as 2.06±0.03, 5.32±1.08, 3.42±1.48, and 3.54±1.24% respectively for concentrations 50, 100, 150, and 200 µg mL^-1^. At 4 h AQPEF treatment and 4 h T+16 h RS, the polar deviation frequency consequently decreased as compared to 2 h treated samples (Table 1, Figure 1). Colchicine (150 µg mL^-1^) induced polar deviation 1.00±0.07 and 0.85±0.85% respectively at 4 h AQPEF treatment and 4 h T+16 h RS. The present investigation shows that AQPEF induced increased polar deviation frequencies at 2 and 4 h treatments but it reduced in 4 h T+16 h RS indicating its reversible effects like colchicine. The polar deviation is a type of mitotic abnormality that was reported earlier with LAECV and colchicine in *A. cepa* root apical meristem cells (Kundu *et al*. 2016). The polar deviation becomes evident in microtubule destabilizing drugs i.e., colchicine. Thus, the preliminary mode of action of AQPEF is almost certainly as reminiscent of colchicine.

An increased percentage of the vagrant chromosomes were observed in the case of 4 h T+16 h RS than 2 and 4 h of AQPEF treatments. The frequency of vagrant chromosome was scored to be 3.69±0.91, 5.16±0.68, and 5.04±0.83 % respectively for the concentration of 100, 150, and 200 µg mL^-1^ of AQPEF in 16 h recovery samples (Table 1). The AQPEF treatment induced an increased percentage of cells with laggard chromosomes at early hours (2 h) of treatment. Here, 50, 100, 150, and 200 µg mL^-1^ concentration of AQPEF induced respectively 1.03±0.60, 2.29±0.80, 0.40±0.40, and 0.81±0.41% of laggard chromosome containing cells at 2 h of treatment (Table 1, Figure 1). Mitotic abnormalities like vagrant chromosomes and laggard chromosomes can also form due to spindle poisoning. Here, in *A. cepa* root tip cells, AQPEF induced both types of abnormalities at 2 and 4 h of treatment in *A. cepa* root tip cells. The frequencies of vagrant and laggard chromosomes were decreased in 4 h of AQPEF treatment followed by 16 h recovery, indicating the induced effects are reversible like colchicine action.

The AQPEF treatment induced a significant (p<0.0001) increase in micronuclei frequency and were scored 4.31±0.33, 5.08±0.13, 5.05±0.22&3.05±0.37% respectively for the concentrations 50, 100, 150, and 200 µg mL^-1^. However, the incidence of micronuclei was not observed in the case of early hours (2 h and 4 h) of AQPEF treated samples (Table 1, Figure 1). Above result correlates with the result obtained from colchicine treatment, where 150 µg mL^-1^ colchicine induces 6.11±0.43% micronucleus in 4 h T+16 h RS. Data indicates that the AQPEF and colchicine treatments on onion root tip cells could induce a significantly (p<0.0001) increased polyploid cells frequency. The polyploidy was induced only in the case of 4 h treatment followed by 16 h water recovery samples in both AQPEF and colchicine treatment. The highest polyploid frequency (20.14±0.68 %) was found in 100 µg mL^-1^ followed by 150 µg mL^-1^ (18.43±0.21 %) and 200 µg mL^-1^ (12.42±0.81 %) concentrations of AQPEF treatment (Table 1, Figure 1). 150 µg m^L-^1 colchicine treatment induces 16.03±0.20% polyploidy. Here, 150 µg mL^-1^ is the most effective concentration of AQPEF to decipher its activity similar to colchicine (150 µg mL^-1^). Disruption of the mitotic spindle inhibits cytokinesis and such inhibition of cytokinesis and reconstruction of nuclei leads to the formation of polyploid cells (Carvalho *et al*. 2019, Fenech and Crott 2002, Chauhan and Sundararaman1990, Chauhan *et al*. 1986, Levan *et al*. 1938). In addition to polyploid cells, microscopic analysis also revealed that AQPEF induces micronuclei in *A. cepa* root apical meristem cells at 4 h T+16 h RS. Reconstruction of c-metaphases, vagrant chromosomes, and chromatid breaks result in the formation of micronuclei. A positive correlation between the formation of c-metaphase, vagrant chromosome, and polyploidy was evident in *A. cepa* root tip cells (Carvalho *et al*. 2019, Levan *et al*. 1938). In conclusion, petroleum ether is suitable solvent that can extract the bioactive phytochemicals of LAECV having cytotoxic effect on *A. cepa* root tip cells. The AQPEF of LAECV has shown to have effective micronuclei, polyploidy, and mitotic abnormality inducing potentials in *A. cepa* root apical meristem cells.

## Acknowledgements

The authors acknowledge Prof. A. Mukherjee for plant species authentication and the financial support of UGC-SRF (FC(Sc)/RS/UGC/ZOO/2018-19/129, w.e.f. 07.04.2018, dated: 04.02.2019), and the DST-PURSE, DST-FIST, and UGC-DRS and MRP sponsored facilities in the Department of Zoology.

## References

AleyKutty, N. A., Mathews, S. M. and Leena, P. N. 2011. Physical, Phytochemical Screening and Antibacterial Activities of Clerodendrum infortunatum L. Root. Int J Pharma Bio Sci. 2(2): 182–187.

Arvind, K., Pradeep, R., Deepa, R. and Mohan, V. 2002. Diabetes and coronary artery diseases. Indian J Med Res.116: 163–176.

Azad, A. K., Wan –Azizi, W. S., Syafiq, T. M. F., Mahmood, S., Al moustafa, H. A. and Labu, Z. K. 2013. Isolation, characterization and cytotoxic effect exploration of methanolic extract of local medicinal plant Clerodendrum viscosum Vent. AJBAS. 7: 641–647.

Babich, H., Segall, M. A. and Fox, K. D. 1997. The Allium test – A simple, eukaryote genotoxicity Assay. Am Biol Teach. 59(9): 580–583

Bakare, A. A., Mosuro, A. A. and Osibanjo, O. 2000. Effect of simulated leachate on chromosomes and mitosis in roots of Allium cepa (L). J. Environ. Biol. 21(3): 263– 271.

Bhattacharjee, D., Das, A., Das, SK. and Chakraborthy, G.S. 2011. Clerodendrum infortunatum L.: A review. J Adv Pharm Healthcare Res. 1: 82–5.

Caritá, R. and Marin-Morales, M. A. 2008. Induction of chromosome aberrations in the Allium cepa test system caused by the exposure of seeds to industrial effluents contaminated with azo dyes. Chemosphere. 72(5): 722–5.

Carvalho, M. S., Andrade-Vieira, L. F., dos Santos, F. E., Correa, F. F., das Graças Cardoso, M. and Vilela, L. R. 2019. Allelopathic potential and phytochemical screening of ethanolic extracts from five species of Amaranthus spp. in the plant model Lactuca sativa. Sci. Hortic. 9(245): 90–8.

Chandrashekar, R., and Rao, S. N. 2012. Acute central and peripheral analgesic activity of ethanolic extract of the leaves of Clerodendrum viscosum in rodent models. J. drug deliv. ther.2(5): 105–108.

Chaudhuri, A. and Ray, S. 2015. Antiproliferative activity of phytochemicals present in aerial parts aqueous extract of Ampelocissus latifolia (Roxb.) Planch. on apical meristem cells. Int J Pharma Bio Sci. 6(2): 99–108.

Chauhan, L. K. and Sundararaman, V. 1990. Effect of substituted ureas on plant cells I. Cytological effects of isoproturon on the root meristem cells of Allium cepa. Cytologia. 55(1): 91–8.

Chauhan, L. K. S., Dikshith, T. S. S. and Sundararaman, V. 1986. Effect of deltamethrin on plant cells. I. Cytological effects on the root meristem cells of Allium cepa. Mutat. Res.171: 25–30.

Das, S., Bhattacharya, S., Prasanna, A., Kumar, RBS., Pramanik, G. and Halder, PK. 2011. Preclinical evaluation of antihyperglycemic activity of Clerodendrum infortunatum leaf against streptozotocin-induced diabetic rats. Diabetes ther. 2(2): 92–100.

Devi, O. I., Dutta, B. K. and Choudhury, P. 2013. Allelopathy effect of aqueous extract of Clerodendrum viscosum Vent, Ageratum conyzoides and Parthenium hysterophorus on the seed germination and seedling vigour of chickpea seeds (Cicer arietinum L) in vitro. J. Nat. Appl. Sci.5(1): 37–40.

Fenech, M. and Crott, J.W. 2002. Micronuclei, nucleoplasmic bridges and nuclear buds induced in folic acid deficient human lymphocytes-evidence for breakage-fusion-bridge cycles in the cytokinesis-block micronucleus assay. Mutat. Res. 504(1-2): 131–6.

Ford, J. H. and Correll, A. T. 1992. Chromosome errors at mitotic anaphase. Genome. 35: 702–705.

Gouthamchandra, K., Mahmood, R. and Manjunatha H. 2010. Free radical scavenging, antioxidant enzymes and wound healing activities of leaves extracts from Clerodendrum infortunatum L. Environ. Toxicol. Pharmacol. 30: 11–18.

Gupta, R. and Singh, H. K. 2012. Nootropic potential of Alternanthera sessilis and Clerodendrum infortunatum leaves on mice. Asian Pac. J. Trop. Dis. 2: 465–470.

Hamilton, F. 1997. The flora homoeopathica. New Delhi: B Jain (Original Publication, 1852).

Hossain, M. S., Islam, J., Sarkar, R. and Hossen, S. M. M. 2014. Antidiarrheal, antidiabetic, antioxidant and antimicrobial activity of methanolic extracts of leaves of Clerodendrum viscosum Vent. Int. j. pharmacogn.1(7): 449–53.

Islam, Md S., Moghal, Md MR., Ahamed, SK., Ahmed, J. and Islam, Md A. 2013. A study on cytotoxic and anthelmintic activities of crude extracts of leaves of Clerodendrum viscosum. Int. Res. J. Pharm. 4: 99–102.

Jirovetz, L., Buchbauer, G., Puschmann, C., Shafi, M. P. and Saidutty, A. 1999. Essential Oil Analysis of the Leaves and the Root Bark of the Plant Clerodendrum infortunatum used in Ayurvedic Medicine. Herba Pol. 45: 87–93.

Kirtikar, K.R. and Basu, B. D. 1991. Indian Medicinal Plants.III.2nd ed. Dehradun: Bishen Singh and Mahendra Pal Singh.

Kundu, L. M. and Ray, S. 2016. Mitotic abnormalities and micronuclei inducing potentials of colchicine and leaf aqueous extracts of Clerodendrum viscosum Vent. in Allium cepa root apical meristem cells. Caryologia. 70: 1(7-14)

Levan, A. 1938. The effect of Colchicine on root mitoses in Allium. Hereditas.24(4): 471–86.

Muh, T., Waliullah, A., Yeasmin, A. M., Wahedul, I. M. and Parvez, H. 2014. Insecticidal and Repellent Activity of Clerodendrum viscosum Vent. (Verbenaceae) Against Tribolium castaneum (Herbst) (Coleoptera: tenebrionoidea). Acad. J. Entomol.7(2): 63–69.

Nadkarni, K. M. and Nadkarni, A. K. 2002. Indian Materia Medica. Bombay: Popular.

Nandi, S. and Lyndem, L. M. 2015. Clerodendrum viscosum: traditional uses, pharmacological activities and phytochemical constituents. Nat. Prod. Res. 30(5): 497–506.

Qasem, J. R. and Foy, C. L. 2001. Weed Allelopathy, Its Ecological Impacts and Future Prospects. J Crop Prod. 4(2): 43–119.

Ray, S., Kundu, L. M., Goswami, S. and Chakrabarti, C. S. 2012. Antiproliferative and apoptosis inducing activity of allelochemicals present in leaf aqueous extract of traditionally used antitumor medicinal plant, Clerodendrum viscosum Vent. IJPRD. 4(06): 332–345.

Ray, S., Kundu, L. M., Goswami, S., Roy, G. C., Chatterjee, S., Dutta, S., Chaudhuri, A. and Chakrabarti, C. S. 2013. Metaphase arrest and delay in cell cycle kinetics of root apical meristems and mouse bone marrow cells treated with leaf aqueous extract of Clerodendrum viscosum Vent. Cell Prolif. 46: 109–117.

Roy, G. C. and Ray, S. 2017. Antiproliferative and apoptosis inducing activities of Leaf organic solvent extract fractions of Clerodendrum viscosum Vent.Int J Pharma Bio Sci. 8(3): 58–66.

Roy, A. and Roy, S. 2019. Assessment of cytotoxic effects of aqueous and methanolic leaf extracts of Clerodendrum inerme (L.) Gaertn. and C. viscosum Vent. using Allium test. Cytologia 84: 73–76.

Sharma, A. K., and Sharma, A. 1999. Plant chromosomes: Analysis, manipulation and engineering. Hardwood Academic Publishers, Netherlands.

Singh, V. K., Ali, Z. A. and Siddiqui, M. K. 1997. Medicinal plants used by the forest ethnics of Gorakhpur district (Uttar Pradesh), India. Int. J. Pharm. 35: 194–206.

Singhmura, Saroj. 2016. A comprehensive overview of a traditional medicinal herb: Clerodendrum infortunatum Linn. J. pharm. sci. 5: 80–84.

Warrier, P. K., Nambbiar, V. P. K. and Kutty, R. C. 1996. In: Indian Medicinal Plants, Orient Longman, Arya Vaidyasala Publication, Hyderabad, India. 1: 160.

Yusuf, M., Chowdhury, J. U., Wahab, M. A. and Begum, J. A. 1994. Medicinal plants of Bangladesh. Dhaka. Bangladesh J Sci Ind Res. 1–340

